# Further antibody escape by Omicron BA.4 and BA.5 from vaccine and BA.1 serum

**DOI:** 10.1101/2022.05.21.492554

**Authors:** Aekkachai Tuekprakhon, Jiandong Huo, Rungtiwa Nutalai, Aiste Dijokaite-Guraliuc, Daming Zhou, Helen M. Ginn, Muneeswaran Selvaraj, Chang Liu, Alexander J. Mentzer, Piyada Supasa, Helen M.E. Duyvesteyn, Raksha Das, Donal Skelly, Thomas G. Ritter, Ali Amini, Sagida Bibi, Sandra Adele, Sile Ann Johnson, Bede Constantinides, Hermione Webster, Nigel Temperton, Paul Klenerman, Eleanor Barnes, Susanna J. Dunachie, Derrick Crook, Andrew J Pollard, Teresa Lambe, Philip Goulder, OPTIC consortium, ISARIC4C consortium, Elizabeth E. Fry, Juthathip Mongkolsapaya, Jingshan Ren, David I. Stuart, Gavin R Screaton

**Author notes:** These authors contributed equally to this work (Aekkachai Tuekprakhon, Jiandong Huo, Rungtiwa Nutalai). See acknowledgements (OPTIC consortium, ISARIC4C consortium).

## Abstract

The Omicron lineage of SARS-CoV-2, first described in November 2021, spread rapidly to become globally dominant and has split into a number of sub-lineages. BA.1 dominated the initial wave but has been replaced by BA.2 in many countries. Recent sequencing from South Africa’s Gauteng region uncovered two new sub-lineages, BA.4 and BA.5 which are taking over locally, driving a new wave. BA.4 and BA.5 contain identical spike sequences and, although closely related to BA.2, contain further mutations in the receptor binding domain of spike. Here, we study the neutralization of BA.4/5 using a range of vaccine and naturally immune serum and panels of monoclonal antibodies. BA.4/5 shows reduced neutralization by serum from triple AstraZeneca or Pfizer vaccinated individuals compared to BA.1 and BA.2. Furthermore, using serum from BA.1 vaccine breakthrough infections there are likewise, significant reductions in the neutralization of BA.4/5, raising the possibility of repeat Omicron infections.

## Introduction

SARS-CoV-2 emerged in Wuhan in late 2019 to rapidly cause a global pandemic. It is now estimated to have infected over half a billion people and caused over 6 million deaths (https://covid19.who.int/). Being a positive sense RNA virus, SARS-CoV-2 was predicted to mutate and that has indeed been the case. Because of the scale of the pandemic it is estimated all single point mutations in the large SARS-CoV-2 genome will be generated every day (Sender et al., 2021). Most mutations will be silent, deleterious or of little consequence, however a few may give the virus an advantage leading to rapid natural selection (Domingo, 2010). Many thousands of individual mutations have been described, and about a year after the outbreak started, strains began to emerge containing multiple mutations particularly in the spike (S) gene. Several of these have been designated variants of concern (VoC) (https://www.cdc.gov/coronavirus/2019-ncov/variants/variant-classifications.html) and have led to successive waves of infection: first Alpha (Supasa et al., 2021), then Delta (Liu et al., 2021a), then Omicron (Dejnirattisai et al., 2022) spread globally becoming the dominant variants. Alongside these, Beta (Zhou et al., 2021) and Gamma (Dejnirattisai et al., 2021b) caused large regional outbreaks in Southern Africa and South America respectively but did not dominate globally. As of 29^th^ April, over 2.5 million cases of Omicron (BA.1 and BA.2) have been reported in the UK alone (https://www.gov.uk/government/publications/covid-19-variants-genomically-confirmed-case-numbers/variants-distribution-of-case-data-29-april-2022#omicron) and, although the disease is less severe, particularly in the vaccinated, the scale of the outbreak has still led to a large number of deaths (Nealon and Cowling, 2022).

S is the major surface glycoprotein on SARS-CoV-2 and assembles into extended transmembrane anchored trimers (Walls et al., 2020; Wrapp et al., 2020) which give virions their characteristic spiky shape. S is divided into N-terminal S1 and C-terminal S2 regions. S1 contains the N-terminal domain (NTD) and receptor binding domain (RBD). A small 25 amino acid (aa) patch at the tip of the RBD is responsible for interaction with the cellular receptor ACE2 (Lan et al., 2020). Following ACE2 binding, S1 is cleaved and detaches, whilst S2 undergoes a major conformational change to expose the fusion loop, which mediates fusion of viral and host membranes, allowing the viral RNA to enter the host cell cytoplasm and commence the replicative cycle (Walls et al., 2017).

S is the major target for neutralising antibodies, and studies by a number of groups have isolated panels of monoclonal antibodies from infected or vaccinated volunteers (Barnes et al., 2020; Dejnirattisai et al., 2021a; Yuan et al., 2020a). Potently neutralizing antibodies are largely confined to three sets of sites on S1. The first is within the NTD (Cerutti et al., 2021; Chi et al., 2020), these antibodies do not block ACE2 interaction and their mechanism of action is still not well determined. A second region of binding is on or in close proximity to the ACE2 binding surface of the RBD; most potently neutralizing antibodies bind this region and prevent interaction of S with ACE2 on the host cell, blocking infection (Dejnirattisai et al., 2021a; Yuan et al., 2020a). Finally, some potent antibodies bind the RBD but do not block ACE2 binding, exemplified by mAb S309 which binds in the region of the N-linked glycan at position 343 (Pinto et al., 2020), these antibodies may function to destabilize the S-trimer (Huo et al., 2020b; Yuan et al., 2020b; Zhou et al., 2020).

Although mutations in the VoC are spread throughout S, there are particular hotspots in the NTD and RBD, exactly where potent neutralizing antibodies bind and they are likely being driven by escape from the antibody response following natural infection or vaccination. Mutation of the ACE2 interacting surface may also give advantage by increased ACE2 affinity for S, or possibly altering receptor tropism (Zahradnik et al., 2021). Increased ACE2 affinity has been found in VoC compared to ancestral strains (Dejnirattisai et al., 2021b; Liu et al., 2021a; Supasa et al., 2021; Zhou et al., 2021), potentially conferring a transmission advantage, but affinity is not increased in Omicron BA.1 (Dejnirattisai et al., 2022) and only marginally in BA.2 (Nutalai et al., 2022).

The initial Omicron wave was caused by the BA.1 strain which, compared to ancestral strains, contains 30 aa substitutions, 6 aa deletions and 3 aa insertions, largely clustered at the sites of interaction of potently neutralizing antibodies: the ACE2 interacting surface; around the N-343 glycan, and in the NTD (Dejnirattisai et al., 2022). These changes cause large reductions in the neutralization titres of vaccine or naturally immune serum, leading to high-levels of vaccine breakthrough infections and contributing to the huge spike in Omicron infection (Dejnirattisai et al., 2022; McCallum et al., 2022).

A number of Omicron sub-lineages have been described. BA.2 and BA.3 were reported at about the same time as BA.1 and are highly related, but contain some unique changes in S (Figure 1A), whilst another sub-lineage BA.1.1, which contains an additional R346K mutation also emerged (Nutalai et al., 2022). The BA.2 strain, which possesses a small transmission advantage, has become globally dominant. BA.3, reported in relatively few sequences compared to BA.1 and BA.2, appears to be a mosaic of BA.1 and BA.2 changes (with 3 differences in the RBD compared to BA.1 and 3 differences compared to BA.2). Cases of BA.2 infection following BA.1, are not thought to be common, due to good levels of cross-neutralizing antibody following vaccination (Nutalai et al, 2022, https://www.who.int/news/item/22-02-2022-statement-on-omicron-sublineage-ba.2).

**Figure 1.**
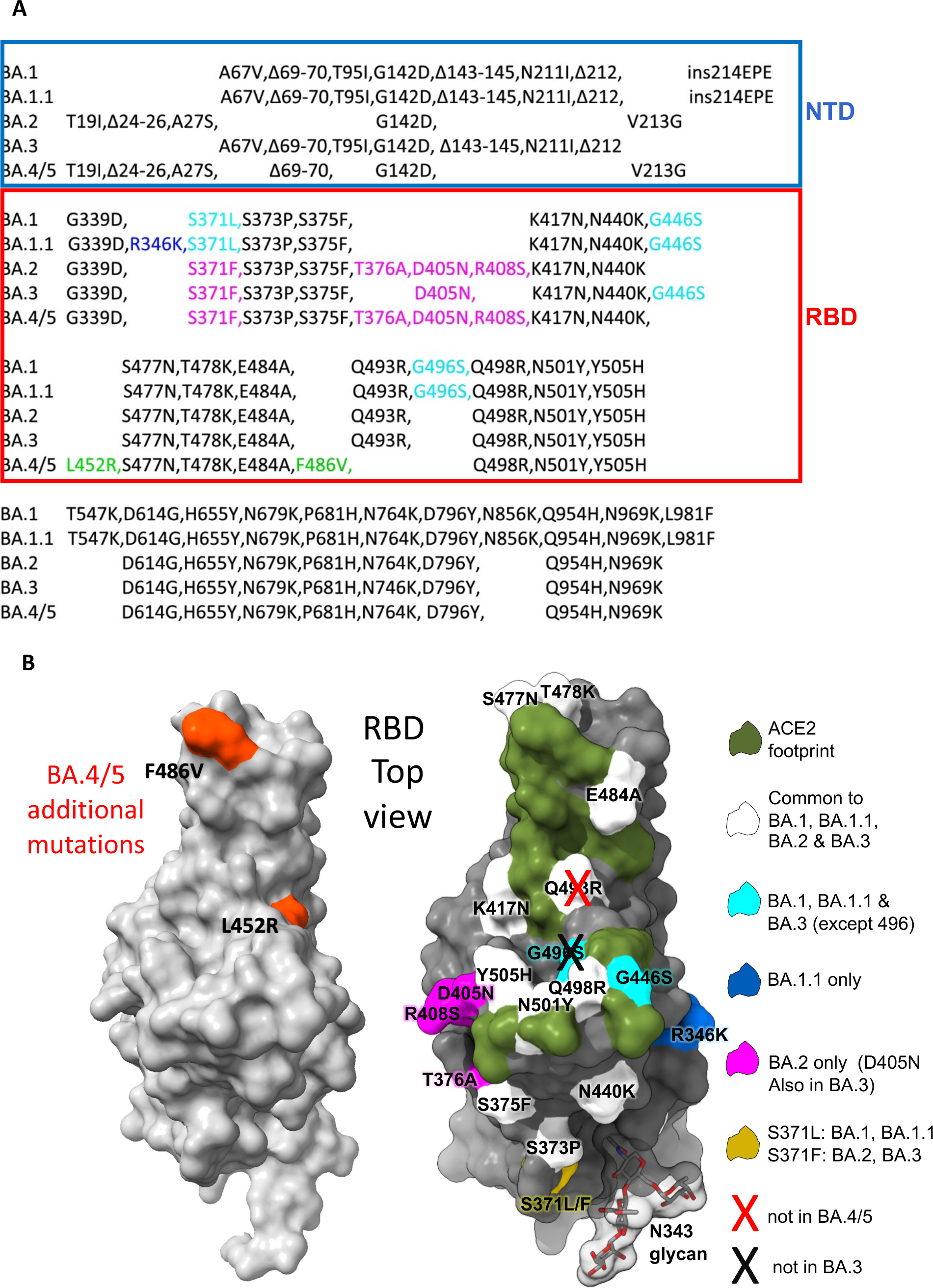
The Omicron sub-lineage compared to BA.4/5. (A) Comparison of S protein mutations of Omicron BA.1, BA.1.1, BA.2, BA.3 and BA.4/5 with NTD and RBD boundaries indicated. (B) Position of RBD mutations (grey surface with the ACE2 footprint in dark green). Mutations common to all Omicron lineages are shown in white (Q493R which is reverted in BA.4/5 is shown with a cross), those common to BA.1 and BA.1.1 in cyan, those unique to BA.1.1 in blue and those unique to BA.2 in magenta. Residue 371 (yellow) is mutated in all Omicron viruses but differs between BA.1 and BA.2. The N343 glycan is shown as sticks with a transparent surface.

In early April 2022 two new Omicron lineages were reported from Gauteng in South Africa and designated BA.4 and BA.5 (https://assets.publishing.service.gov.uk/government/uploads/system/uploads/attachment_data/file/1067672/Technical-Briefing-40-8April2022.pdf). These have become dominant in Gauteng and look to be fuelling a new wave of infection in South Africa, with some international spread. BA.4 and BA.5 (from here on referred to as BA.4/5), have identical S sequences, and appear to have evolved from BA.2. They contain additional mutations in the RBD; in particular the reversion mutation R493Q, together with mutations L452R and F486V (Figure 1A).

Here we report the antigenic characterisation of BA.4/5 compared to the other Omicron sub-lineages (for completeness we also report data on BA.3, although this is of less concern). We find neutralization of BA.4/5 by triple dosed vaccine serum is reduced compared to BA.1 and BA.2. We also see reductions in titres against BA.4/5 compared to BA.1 and BA.2 in sera from cases who had suffered vaccine breakthrough BA.1 infections. Neutralization of the Omicron lineage by a panel of recently derived potent Omicron specific mAbs, raised following vaccine breakthrough BA.1 infection (Nutalai et al., 2022) is reduced: 10/28 are completely knocked out against BA.4/5, while several others suffer large reductions in activity compared to the other Omicron lineages. We corroborate the neutralisation results with biophysical analysis of binding, and provide structure-function explanations for mAb failure against BA.4/5 with the changes at residues 452 and 486, both of which cause serious impact. Finally, we measure the affinity of the BA.4/5 RBD for ACE2 and find that it is higher than ancestral Omicron strains.

## Results

### The Omicron lineages BA.4/5

BA.4 and BA.5 S sequences are identical, and closely related to BA.2 (sequence diversity in Omicron S is shown in Figure 1A). Compared to BA.2, BA.4/5 has residues 69 and 70 deleted, and contains 2 additional substitutions in the RBD: L452R and F486V, finally BA.4/5 lacks the Q493R change seen in BA.1 and BA.2, reverting to Q493 as in the Victoria/Wuhan strain.

The 2 additional mutations in the RBD are of most concern in terms of antibody escape: L452R is a chemically radical change and is one of the pair of changes in Delta RBD (the other, T478K, is already found in the Omicron lineage). Mutation F486L was found in sequences of SARS-CoV-2 isolated from Mink early in the pandemic and is also a site of escape mutations to several mAbs (Gobeil et al., 2021). The change F486V in BA.4/5 is also a reduction in the bulk of the hydrophobic side-chain as in F486L, but more significant. Both residues 452 and 486 lie close to the edge of the ACE2 interaction surface (Figure 1B) and, together with the reversion to ancestral sequence Q493 which lies within the ACE2 footprint, have the potential to modulate ACE2 affinity and the neutralizing capacity of vaccine or naturally acquired serum. The L452R and F486V mutations are likely to cause more antibody escape, while the reversion at 493 may reduce the escape from responses to earlier viruses.

### Neutralization of BA.4/5 by vaccine serum

We constructed a panel of pseudotyped lentiviruses (Di Genova et al., 2020) expressing the S gene from the Omicron sub-lineages BA.1, BA.1.1, BA.2, BA.3 and BA.4/5 together with early pandemic Wuhan related strain, Victoria, used as control. Neutralization assays were performed using serum obtained 28 days following a third dose of the Oxford-AstraZeneca vaccine AZD1222 (n = 41) (Flaxman et al., 2021) or of Pfizer-BioNtech vaccine BNT162b2 (n = 20) (Cele et al., 2021a) (Figure 2 A,B). For AZD1222, neutralization titres for BA.4/5 were reduced 2.1-fold compared to BA.1 (p<0.0001) and 1.8-fold compared to BA.2 (p<0.0001).

**Figure 2.**
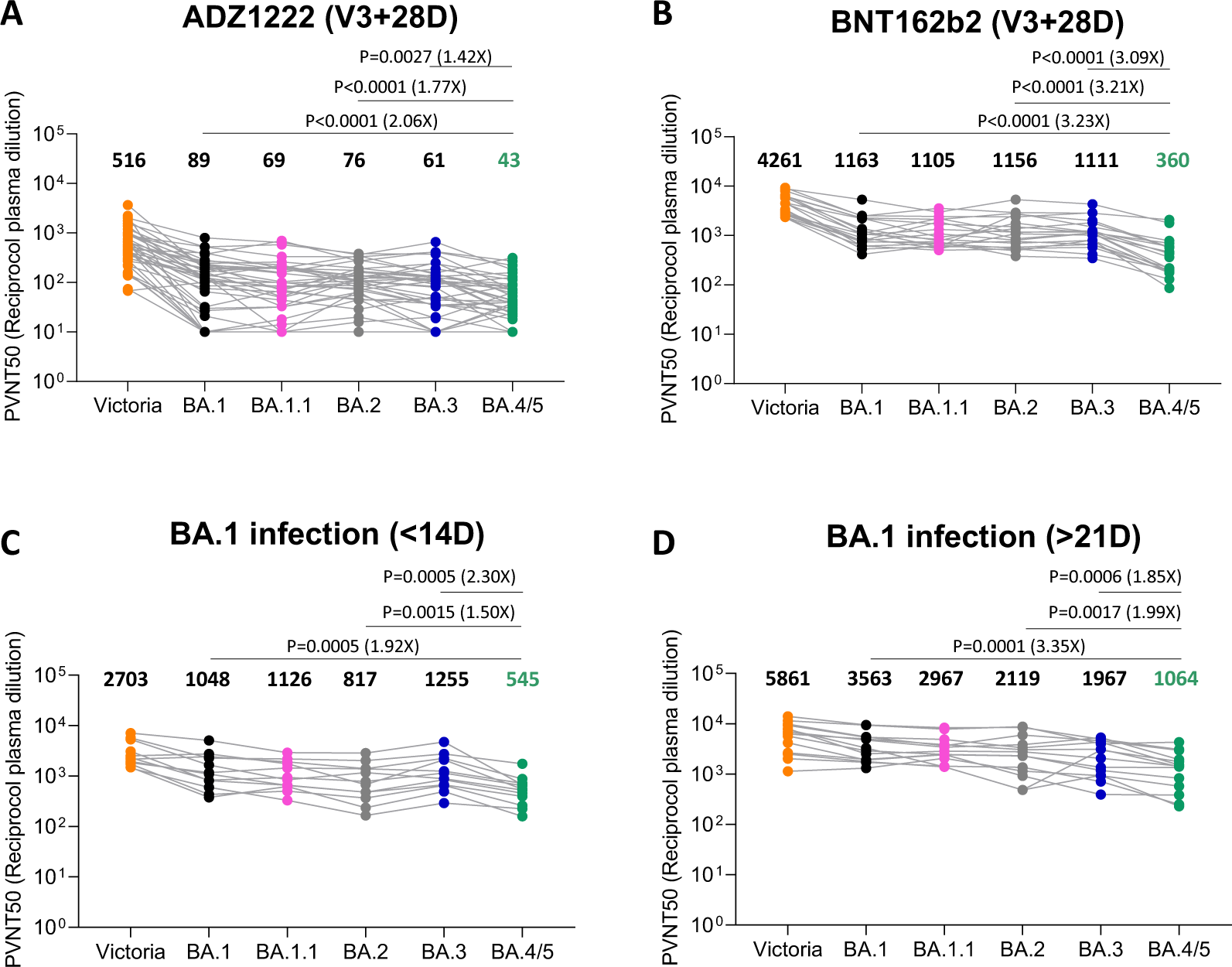
Pseudoviral neutralization assays of BA.4/5 by vaccine and BA.1 immune serum. IC50 values for the indicated viruses using serum obtained from vaccinees 28 days following their third dose of vaccine (A) AstraZeneca AZD AZD1222 (n=41), (B) 4 weeks after the third dose of Pfizer BNT162b2 (n=20). Serum from volunteers suffering breakthrough BA.1 infection volunteer taken (C) early ≤14 (n=12) days from symptom onset (median 13 days) (D) late ≥ 21 days from symptom onset (median 38 days) n=16. Comparison is made with neutralization titres to Victoria an early pandemic strain, BA.1, BA.1..1, BA.2 and BA.3. Geometric mean titres are shown above each column. The Wilcoxon matched-pairs signed rank test was used for the analysis and two-tailed P values were calculated.

For BNT162b2, neutralization titres were reduced 3.2-fold (p<0.0001) and 3.2-fold (p<0.0001) compared to BA.1 and BA.2 respectively. These reductions in titre are likely to reduce vaccine effectiveness at preventing infection, particularly at longer time points as antibody titres naturally wane although it would be expected protection would remain against severe disease.

### Neutralization of BA.4/5 by serum from breakthrough BA.1 infection

Early in the Omicron outbreak we recruited vaccinated volunteers who had all suffered breakthrough Omicron infections. Samples were first taken ≤14 days from symptom onset (median 13 days), while late samples were taken ≥ 21 days from symptom onset (median 38 days) n=16. Pseudoviral neutralization assays were performed against the panel of pseudoviruses described above (Figure 2C,D).

As we have previously described, BA.1 infection following vaccination leads to a broad neutralizing response, with high titres to all the VoC, which is boosted at later time points (Nutalai et al., 2022). Neutralization titres against BA.4/5 were significantly less than BA.1 and BA.2; at the early time point BA.4/5 titres were reduced 1.9-fold (p=0.0005) and 1.5- fold (p=0.0015) compared to BA.1 and BA.2 respectively. At the later point BA.4/5 titres were reduced 3.4-fold (p=0.0001) and 2-fold (p=0.0017) compared to BA.1 and BA.2 respectively.

Thus, BA.4/5 shows a degree of immune escape from the vaccine/BA.1 response when compared with BA.1 and BA.2. These samples were all taken reasonably close to the time of infection meaning that further waning in the intervening months may render individuals susceptible to reinfection with BA.4/5

### Escape from monoclonal antibodies by BA.4/5

We have recently reported a panel of potent human mAb generated from cases of Omicron breakthrough infection (Nutalai et al., 2022). For the 28 most potent mAbs (BA.1 IC50 titres <100 ng/ml) we used pseudoviral assays to compare BA.4/5 neutralization with neutralization of BA.1, BA.1.1, BA.2 and BA.3 (Figure 3A, Table S1A). Neutralization of BA.4/5 was completely knocked out for 10/28 mAbs. Four further mAbs (Omi-09, 12, 29 and 35) showed >5-fold reduction in the neutralization titre of BA.4/5 compared to BA.2. All of these antibodies interact with the RBD, with the exception of Omi-41, which binds the NTD and specifically neutralizes BA.1, BA.1.1 and BA.3 but not BA.2 or BA.4/5 (for unknown reasons Omi-41 can neutralize WT Victoria virus but not Victoria pseudovirus)(Nutalai et al., 2022).

**Figure 3.**
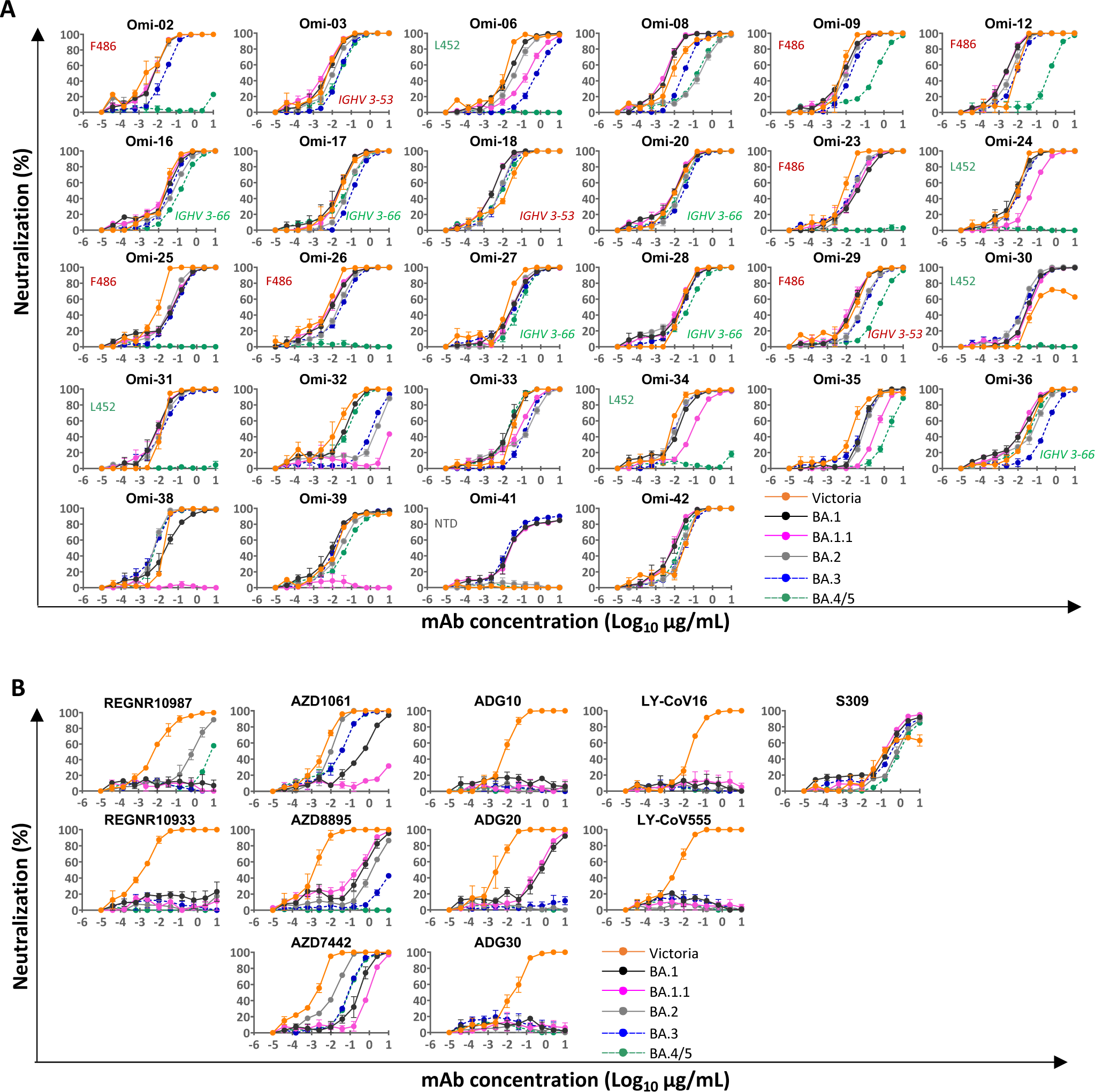
Pseudoviral neutralization assays against Omicron and commercial monoclonal antibodies. (A) Neutralization curves for a panel of 28 monoclonal antibodies made from samples taken from vaccinees infected with BA.1. Titration curves for BA.4/5 are compared with Victoria, BA.1, BA.1.1, BA.2 and BA.3, mAbs we propose to be affected by the L452R and F486L mutations are indicated as are those belonging to the IGVH3-53/66 gene families. (B) Pseudoviral neutralization assays with mAb developed for human use. IC50 titres for mAb in A and B are shown in Table S1.

Sensitivity to L452R: We have previously reported that Omi-24, 30, 31, 34 and 41 show complete knock out of neutralizing activity against Delta, with Omi-06 showing severe knock-down of activity (Nutalai et al., 2022). Since BA.1 and BA.2 harbour only one (T478K) of the 2 Delta RBD mutations, whilst BA.4/5 also harbour L452R, we would expect all five of these L452 directed mAbs to be knocked out on BA.4/5. This is indeed observed (Figure 3A, Table S1A). Omi-41 also fails to neutralize, which is attributed to the differences in mutations in the NTD (Figure 1A).

To confirm that the neutralization effects observed are directly attributable to alterations in RBD interactions we also performed binding analyses of selected antibodies to BA.4/5 and BA.2 RBDs by surface plasmon resonance (SPR) (Figures 4, S2). Omi-31 was chosen as representative of the set of L452R sensitive antibodies, and as expected the binding is severely affected (Figure 4A). Since we have detailed information on the interaction of several Omicron responsive antibodies with the RBD, including Omi-31, we modelled the BA.4/5 RBD mutations in the context of known structures for Omicron Fabs complexed with BA.1 or Delta RBDs (Dejnirattisai et al., 2022; Nutalai et al., 2022), (Figure 5). The Omi-31 complex is shown in Figure 5A and shows L452 tucked neatly into a hydrophobic pocket, which is unable to accommodate the larger positively charged arginine in BA.4/5 and Delta.

**Figure 4.**
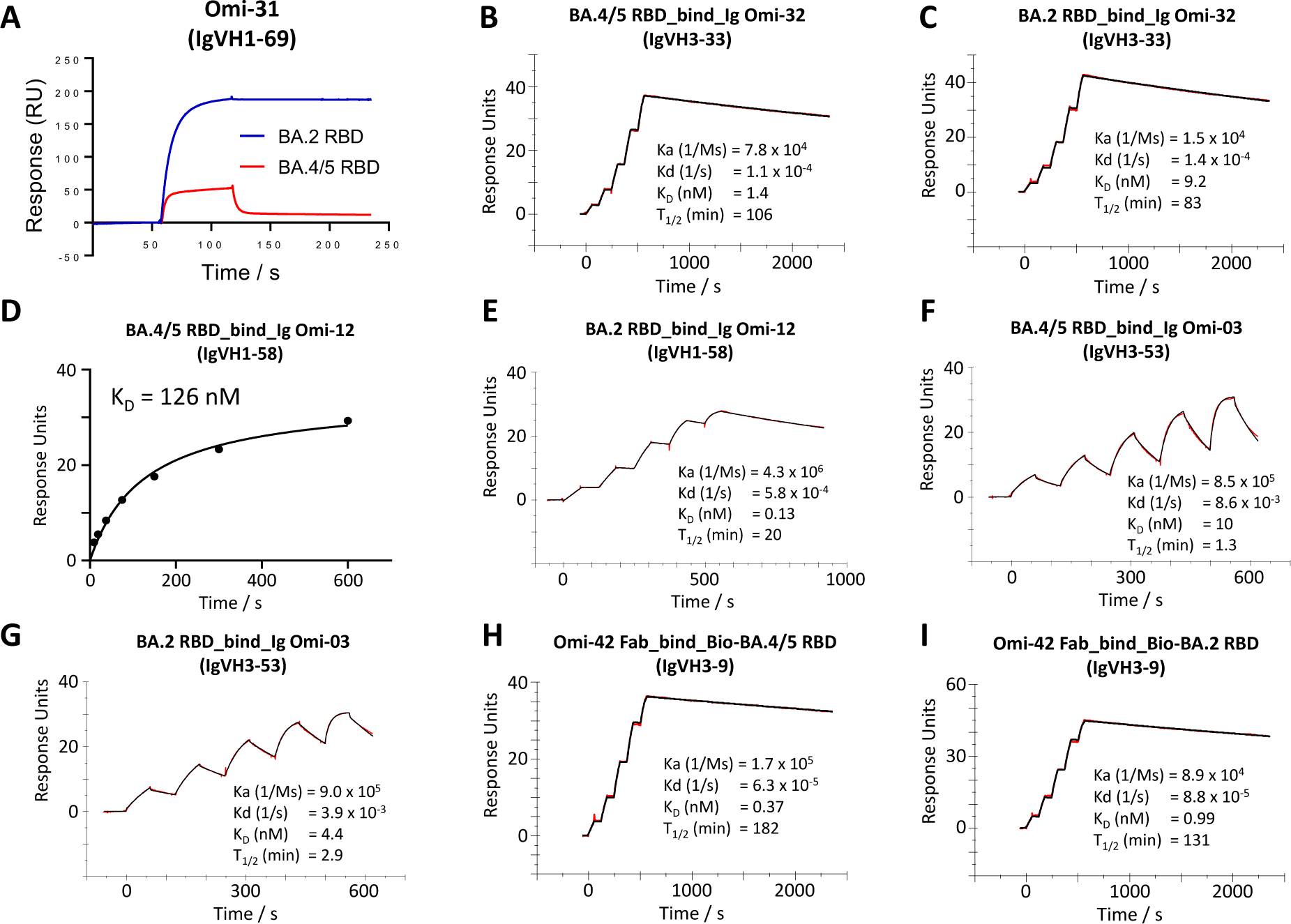
Surface plasmon resonance (SPR) analysis of interaction between BA.2 or BA.4/5 RBD and selected mAbs. (A) Binding of BA.4/5 RBD is severely reduced compared to that of BA.2, so that the binding could not be accurately determined, as shown by a single-injection of 200 nM RBD over sample flow cells containing IgG Omi-31. (B-C; E-I) Sensorgrams (Red: original binding curve; black: fitted curve) showing the interactions between BA.2 or BA.4/5 RBD and selected mAbs, with kinetics data shown. (D) Determination of the affinity of BA.4/5 RBD to Omi-12 using a 1:1 binding equilibrium analysis.

**Figure 5.**
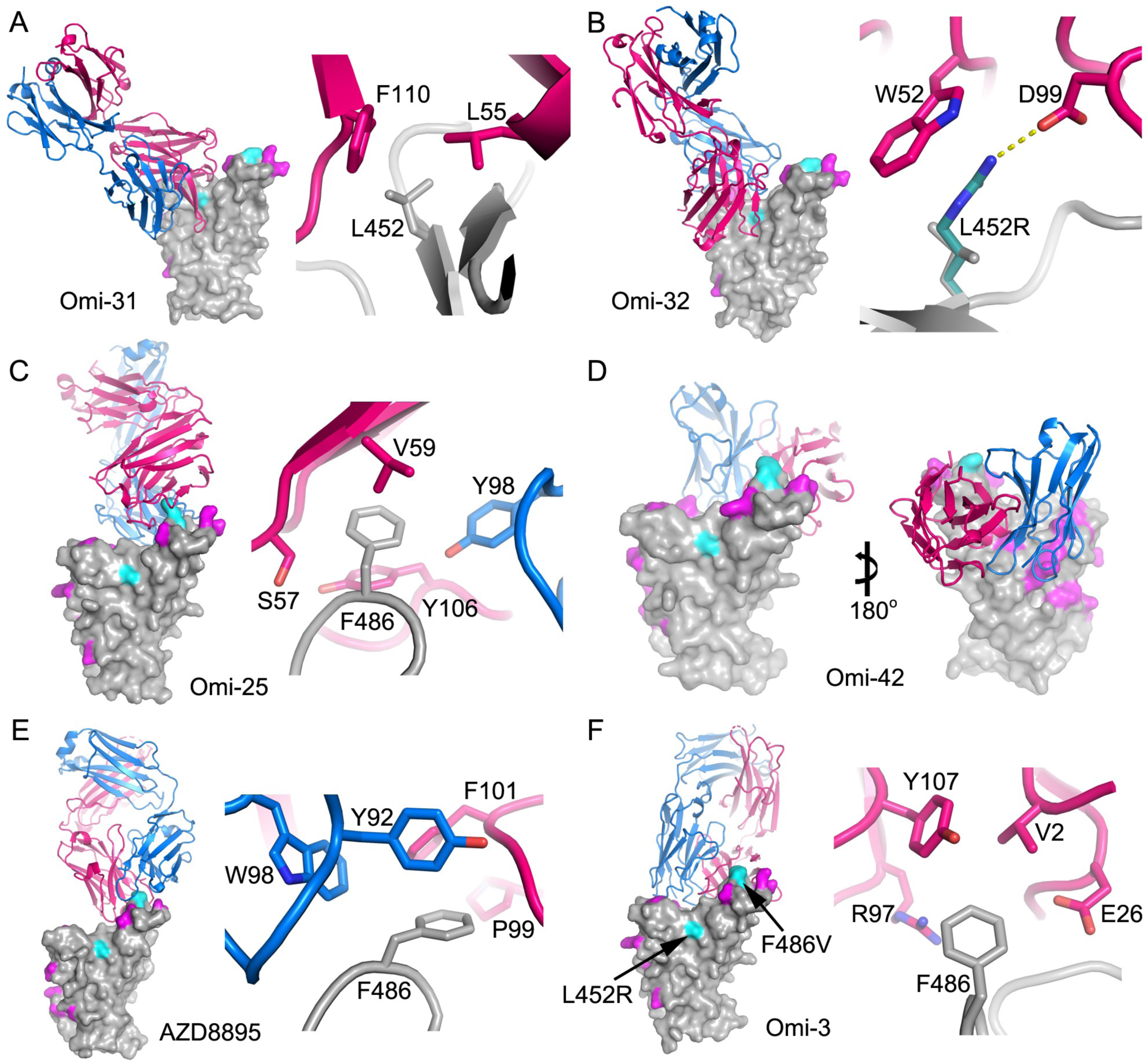
Interactions between mAb and BA.4/5 mutation sites. Overall structure (left panel) and interactions (≤ 4 Å) with BA.4/5 mutation sites (right panel) for (A) BA.1- RBD/Omi-31 (PDB 7ZFB), (B) BA.1-RBD/Omi-32 (PDB 7ZFE), (C) BA.1-RBD/Omi-25 (PDB 7ZFD), (D) BA.1-RBD/Omi-42 (PDB7ZR7), (E) Wuhan-RBD/AZD8895 (PDB 7L7D) and (F) BA.1-RBD/Omi-3 (PDB 7ZF3) complexes. In the left panels RBD is shown as surface representation, with BA.4/5 mutation sites highlighted in magenta and the additional two mutation sites of BA.4/5 at 452 and 486 in cyan, and Fab LC as blue and HC as red ribbons. In the right panel, side chains of RBD, Fab HC and LC are drawn as grey, red and blue sticks, respectively. In (B) L452R (green sticks) are modelled to show a salt bridge to D99 of CDR-H3 may be formed (yellow broken sticks). (D) Beta-RBD/Omi-42 complex showing the Fab does not contact any of the two BA.4/5 mutation sites.

### L452R enhancement of binding

Omi-32 shows 77-fold enhanced neutralization of BA.4/5 compared to BA.2. Kinetic analysis of Fab binding to the RBDs suggests that this is mainly achieved by a 5-fold increase in the on-rate of binding (Figure 4B, C). This is largely explained by the favorable interaction of the arginine at 452 making a salt bridge to residue 99 of the heavy chain (HC) CDR3 (Figure 5B), perhaps assisted by removal of slightly unfavourable charge interactions at residue 493. It is possible that these electrostatic changes enhance on-rate by electrostatic steering of the incoming antibody.

### Sensitivity to F486V

Extending the logic used to understand Delta sensitivity, the remaining antibodies affected by BA.4/5 > BA.2, but which retain activity against Delta, namely Omi- 02, 09, 12, 23, 25, 26, 29, are likely sensitive to the F486V change. The binding sensitivity was confirmed by SPR analysis of Omi-12 (Figure 4D, E) which showed an almost 1,000-fold reduction in affinity. An example of the structural basis of sensitivity is provided by the Omi-25 complex (Figure 5C), which shows that the phenylalanine side chain acts as a binding hot- spot, nestled in a hydrophobic cavity making favorable ring-stacking interactions with Y106 of the HC CDR3.

### Activity of commercial antibodies against BA.4/5

We tested a panel of antibodies that have been developed for therapeutic/prophylactic use against BA.4/5 (Figure 3B, Table S1B). Many of these antibodies have already suffered severe reductions or knock out of activity against BA.1, BA.1.1 or BA.2. For AstraZeneca AZD1061, activity to BA.4/5 was similar to BA.2 (< 2-fold reduction), whilst for AZD8895 residual activity against BA.2 was knocked out. The activity of the combination of both antibodies in AZD7442 (Dong et al., 2021) was reduced 8.1-fold compared with BA.2. The residual activity of REG10987 (Weinreich et al., 2021) against BA.2 was further reduced on BA.4/5, likewise residual BA.1 neutralizing activity was knocked out for ADG20 (Yuan et al., 2022) on BA.4/5. For S309 (VIR-7831/7832) (Sun and Ho, 2020), activity against BA.4/5 was 1.6 fold reduced compared to BA.2.

These effects can be rationalized by reference to the way the antibodies interact with the RBD, for instance in the case of AZD8895 (an IGHV1-58 genotype mAb, Figure 5E), F486 forms a hydrophobic interaction hotspot which will be abrogated by the mutation to a much smaller valine sidechain. Antibody residues involved in the interactions with F486 are highly conserved among this genotype of mAbs, including Omi-12, 253 and Beta-47 (Nutalai et al., 2022; Dejnirattisai et al., 2021a; Liu et al., 2021b), explaining the severe effect of the F486V mutation on neutralization of these mAbs (Figures 3A, S1).

### Systematic themes in mAb interactions

Both Omi-3 (a representative of the IGVH3-53 gene family) and AZD8895 (IGVH1-58) make contacts with F486. Whilst the F486V mutation has little effect on Omi-3 (Figure 4F, G, 5F), it seriously reduces the neutralization of AZD8895 and other IGVH1-58 mAbs e.g. Omi-12 (Figure 4D, E, 5E). It is notable that whereas the numerous Omi series antibodies belonging to the closely related IGVH3-53 and IGVH3-66 gene families (9/28 in total Figure 3A Table S2) are almost entirely resilient to the BA.4/5 changes, the large majority of antibodies from these gene families elicited against earlier variants are knocked out on BA.1 and BA.2 (Nutalai et al., 2022), consistent with selection of a subset of antibodies by breakthrough Omicron infection that are insensitive to the further BA.4/5 mutations.

The effects on antibodies with broadly similar epitopes can vary dramatically, and this is equally true for antibodies which have 452 or 486 central to their binding footprint. Thus Omi-31 (IGVH1-69) and Omi-32 (IGVH3-33), both bind in front of the right shoulder with their CDR-H3 positioned close to 452, whilst the activity of Omi-31 is abolished by L452R (as detailed above), Omi-32 is markedly enhanced (Figure 3A, 5A, B). Similarly, Omi-25 and Omi-42 both belong to the IGVH3-9 gene family and their footprints are in the 486 region (Figure 5C, D). Omi-25 contacts F486, thus neutralization of BA.4/5 is abolished. In contrast Omi-42 does not contact either of the mutation sites and neutralization is fully retained for BA.4/5 (Figure 4H, I, 5D).

### ACE2 RBD affinity

We measured the affinity of BA.4/5 RBD for ACE2 by SPR (Figure 6A-D). The affinity of BA.4/5 RBD was increased compared to the ancestral virus (Wuhan), BA.1 and BA.2 (approximately 3-fold, 3-fold and 2-fold, respectively (BA.4/5/ACE2 KD = 2.4 nM) (Dejnirattisai et al., 2022; Nutalai et al., 2022), which is mainly attributed to an increase in binding half-life, modelling of the ACE2/RBD complex suggests that the bulk of this effect comes from the electrostatic complemantary between ACE2 and the RBD contributed by the L452R mutation (Figure 6E-G).

**Figure 6.**
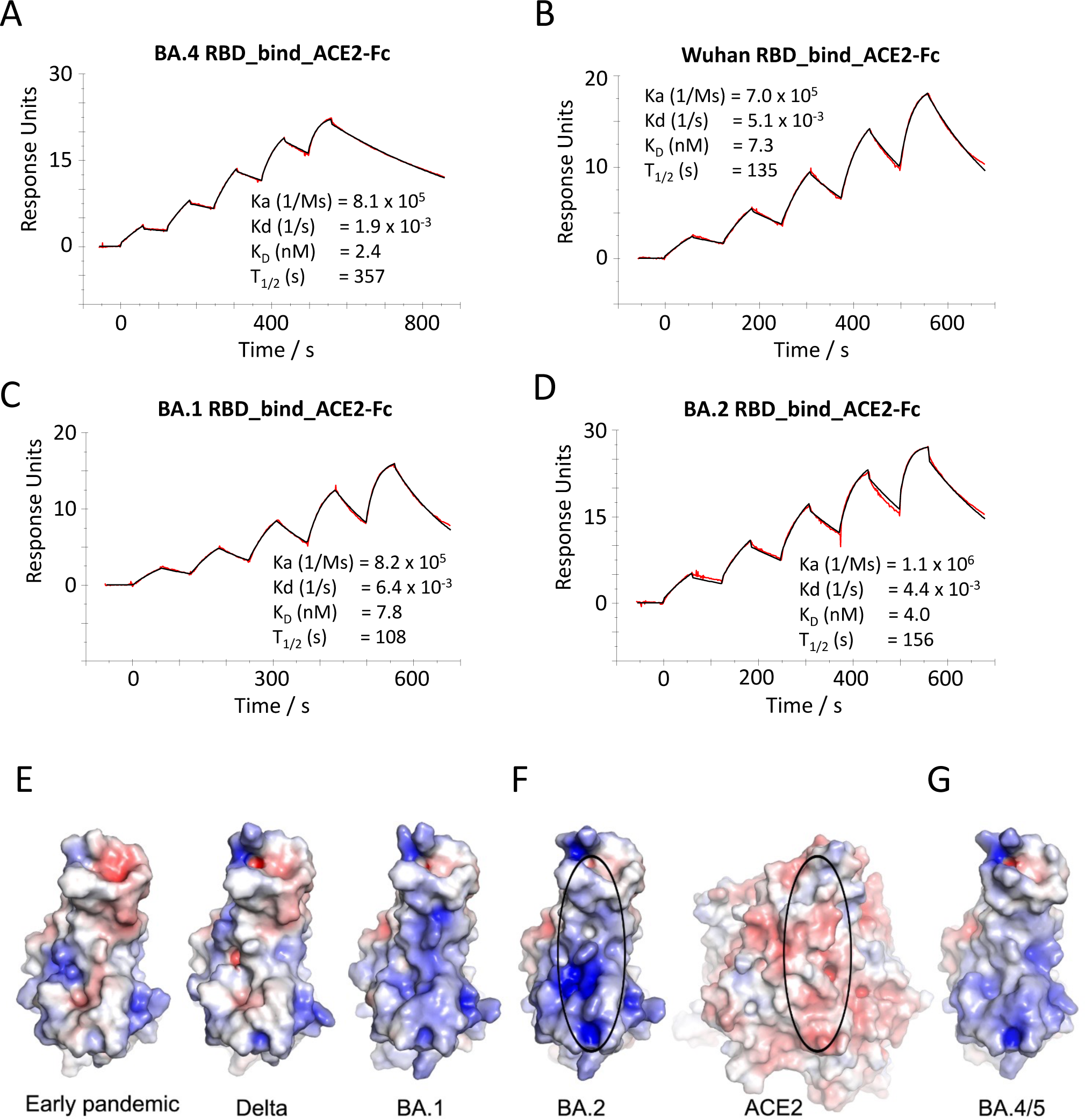
ACE2 RBD affinity. (A)-(D) SPR sensorgrams showing ACE2 binding of BA.4/5 RBD (A) in comparison to binding to ancestral (Wuhan) (B), BA.1 (C) and BA.2 RBD (D). The data for Wuhan, BA.1 and BA.2 have been reported previously in (Nutalai et al., 2022). (E)-(G) Electrostatic surfaces, (E) from left to right, early pandemic, Delta and BA.1 RBD respectively, (F) open book view of BA.2 RBD and ACE2 of the BA.2 RBD/ACE2 complex (PDB 7ZF7), and (G) BA.4/5 RBD (modelled based on the structure of BA.2 RBD).The lozenges on ACE2 and RBD show the interaction areas.

### Antigenic cartography

The neutralization data above has been used to place BA.3 and BA.4/5 on an antigenic map. We repeated the method used for analysis of the Delta and Omicron variants (Liu et al., 2021a), where individual viruses were independently modelled allowing for serum specific scaling of the responses (Methods). The measured and modelled responses are shown in Figure 7A (with 1551 observations and 340 parameters the residual error is 23 %). The results are best visualized in three dimensions, see Video S1, but 2D projections are shown in Figure 7B. This shows, as expected, that the Omicron sub-lineages are clustered together but well separated from early pandemic virus and earlier VoC. Amongst the Omicron cluster BA.4/5 is the most distant from the pre-Omicron viruses.

**Figure 7.**
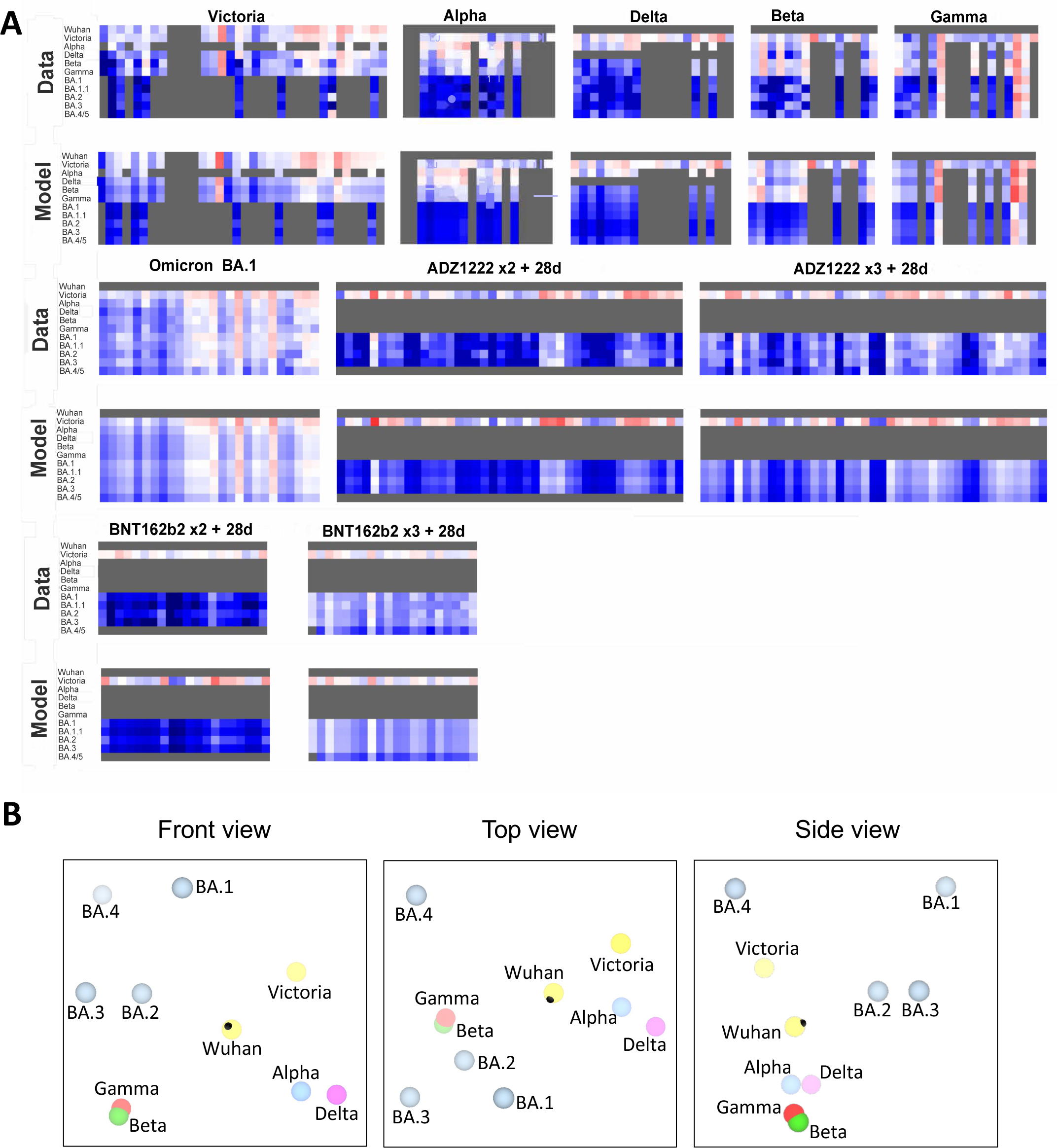
Antigenic mapping. (A) Neutralization data and model (log titre values) used to calculate antigenic maps in (B). Columns represent sera collected from inoculated volunteers or infected patients. Rows are challenge strains: Victoria, Alpha, Delta, Beta, Gamma, BA.1, BA1.1, BA.2, BA.3 and BA.4/5 in order. Values are colored according to their deviation from the reference value; the reference value is calculated on a serum-type basis as the average of neutralization titres from the row which gives this the highest value. (B) Orthogonal views of the antigenic map showing BA.4/5 in the context of the positions of previous VoC and BA.1, BA.1.1, BA.1 and BA.2, calculated from pseudovirus neutralisation data. Distance between two positions is proportional to the reduction in neutralisation titre when one of the corresponding strains is challenged with serum derived by infection by the other.

## Discussion

Following its emergence in November 2019, a succession of SARS-CoV-2 viral variants have appeared with increased fitness, which have rapidly outcompeted the preceding strain and spread globally, the most recent, Omicron appearing in late 2021.

Despite the availability of vaccines, the pandemic has not been brought under control and through Omicron, infections are as high as ever. Although vaccines are effective at preventing severe disease, they are less effective at preventing transmission, particularly of the Omicron sub-lineages. The very high level of viral replication globally drives the accrual of mutations in the viral genome and we are now seeing the assembly of dozens of individual changes in single viruses. Virus recombination, which was predicted, is now being detected, allowing shuffling of complex genomes, such as XD (Delta/BA.1) and XE (BA.1/BA.2), which in the latter case may be more transmissible (https://assets.publishing.service.gov.uk/government/uploads/system/uploads/attachment_data/file/1063424/Tech-Briefing-39-25March2022_FINAL.pdf).

How such large sequence jumps, such as that to the Omicron lineage occur is not known. It has been suggested that these may be occurring in immunocompromised or HIV infected cases, where chronic infections have been documented to last for many months or in some cases over a year. Selection of antibody escape mutations has been documented in such individuals (Cele et al., 2021b; Karim et al., 2021; Kemp et al., 2021) and successive rounds of replication, recombination and perhaps reinfection may be responsible for the selection of the constellation of S mutations found in the Omicron lineage.

BA.4/5, the most recently reported Omicron sub-lineages, seem to be taking hold in South Africa and may spread globally to replace BA.2. Although highly related to BA.2, BA.4/5 contain the 69-70 deletion in the NTD which was also found in Alpha, BA.1 and BA.3, together with additional mutations in the RBD (L452R and F486K). Thus, BA.4/5 has assembled mutations at all of the previously described positions in the VoC Alpha (N501Y), Beta (K417N, E484K, N501Y), Gamma (K417T, E484K, N501Y) and Delta (L452, T478K), the only difference being E484A in BA.4/5 rather than E484K found in Beta and Gamma.

Here, we report greater escape from neutralization of BA.4/5 compared to BA.1 and BA.2. Serum from triple vaccinated donors has ∼2-3-fold reduction in neutralization titres compared to the neutralization of BA.1 and BA.2. Additionally, serum from breakthrough BA.1 infections in vaccinees shows ∼2-3-fold reduction in neutralization titres to BA.4/5 compared to BA.1 and BA.2. These data suggest that a further wave of Omicron infection, driven by BA.4/5 is likely, partly due to breakthrough of vaccine and naturally acquired immunity, although there is no evidence yet of increased disease severity Using a panel of potent mAbs generated from vaccinated cases infected with BA.1 we show the importance of the two new RBD mutations in BA.4/5. The activity of many mAbs is either knocked out or severely impaired against BA.4/5 compared to BA.2. From the neutralization data on BA.4/5, compared to that on Delta, we have been able to impute the contribution of L452R and F486V, and by combining with SPR data, as well as previous mapping by BLI competition matrices and detailed structural data (Nutalai et al., 2022) we are able to understand the basis of these effects on neutralisation and show that the L452R and F486V mutations both make major contributions to BA.4/5 escape.

It is clear that the Omicron lineage, and particularly BA.4/5, has escaped or reduced the activity of mAbs developed for clinical use, although ADZ7442 and S309 still show activity against BA.4/5. New monoclonals and combinations may be needed to plug the gap in activity, to protect the extremely vulnerable and those unable to mount adequate vaccine responses. There is also a question about vaccines, all current vaccines use spike derived from the original virus isolated from Wuhan. Vaccines have been remarkably effective at reducing severe disease and a triple dosing schedule has provided, at least in the short term, protection against Omicron. However, prevention of transmission may become less effective as viruses evolve antigenically further from ancestral strains. Some argue for next-generation vaccines tailored to antigenically distant strains such as Omicron to give better protection, probably used in combination with boosters containing ancestral strains. Whilst vaccination is unlikely to eliminate transmission, the combination of vaccines with boosting by natural infection will probably continue to protect the majority from severe disease.

Finally, it is impossible to say where SARS-CoV-2 evolution will go next, but it is certain the virus will continue to drift antigenically. This may be a continuation along the Omicron lineage or we may see a large jump to a completely new lineage, like the one from Delta to Omicron. The observation that of the 30 aa substitutions in BA.1, all but one was achieved by a single base change in the codon, suggests there remains plenty of antigenic space for SARS-CoV-2 to explore and the capacity for recombination, which has so far not been observed to have breakpoints within the major antigenic sites, could generate more radical antigenic shift.

## Acknowledgements

This work was supported by the Chinese Academy of Medical Sciences (CAMS) Innovation Fund for Medical Science (CIFMS), China (grant number: 2018-I2M-2-002) to D.I.S. and G.R.S. We are also grateful for support from Schmidt Futures, the Red Avenue Foundation and the Oak Foundation. G.R.S. was supported by Wellcome. H.M.E.D., and J.Ren are supported by Wellcome (101122/Z/13/Z), D.I.S. and E.E.F. by the UKRI MRC (MR/N00065X/1). D.I.S. and G.R.S. are Jenner Investigators. This is a contribution from the UK Instruct-ERIC Centre. AJM is an NIHR-supported Academic Clinical Lecturer. The convalescent sampling was supported by the Medical Research Council [grant MC_PC_19059] (awarded to the ISARIC-4C consortium) (with a full contributor list available at https://isaric4c.net/about/authors/) and the National Institutes for Health and Oxford Biomedical Research Centre and an Oxfordshire Health Services Research Committee grant to AJM. OPTIC Consortium: Christopher Conlon, Alexandra Deeks, John Frater, Lisa Frending, Siobhan Gardiner, Anni Jämsén, Katie Jeffery, Tom Malone, Eloise Phillips, Lucy Rothwell, Lizzie Stafford. The Wellcome Centre for Human Genetics is supported by the Wellcome Trust (grant 090532/Z/09/Z). The computational aspects of this research were supported by the Wellcome Trust Core Award Grant Number 203141/Z/16/Z and the NIHR Oxford BRC.

The Oxford Vaccine work was supported by UK Research and Innovation, Coalition for Epidemic Preparedness Innovations, National Institute for Health Research (NIHR), NIHR Oxford Biomedical Research Centre, Thames Valley and South Midland’s NIHR Clinical Research Network. We thank the Oxford Protective T-cell Immunology for COVID-19 (OPTIC) Clinical team for participant sample collection and the Oxford Immunology Network Covid- 19 Response T cell Consortium for laboratory support. We acknowledge the rapid sharing of Victoria, B.1.1.7 and B.1.351 which was isolated by scientists within the National Infection Service at PHE Porton Down, and the B.1.617.2 virus was kindly provided Wendy Barclay and Thushan De Silva. We thank The Secretariat of National Surveillance, Ministry of Health Brazil for assistance in obtaining P.1 samples. This work was supported by the UK Department of Health and Social Care as part of the PITCH (Protective Immunity from T cells to Covid-19 in Health workers) Consortium, the UK Coronavirus Immunology Consortium (UK-CIC) and the Huo Family Foundation. EB and PK are NIHR Senior Investigators and PK is funded by WT109965MA and NIH (U19 I082360). SJD is funded by an NIHR Global Research Professorship (NIHR300791). DS is an NIHR Academic Clinical Fellow. The views expressed in this article are those of the authors and not necessarily those of the National Health Service (NHS), the Department of Health and Social Care (DHSC), the National Institutes for Health Research (NIHR), the Medical Research Council (MRC) or Public Health, England.

## Author Information

These authors contributed equally: A.T., J.H., R.N.

## Contributions

J.H. performed interaction affinity analyses. D.Z. performed antibody competition analyses. D.Z., J.H., J.R., N.G.P., M.A.W., and D.R.H. prepared the crystals and enabled and performed X-ray data collection. J.R., E.E.F., H.M.E.D. and D.I.S. analyzed the structural results. G.R.S., J.H., J.M., P.S., D.Z., R.N., A.T., A.D-G., M.S., R.D. and C.L. prepared the RBDs, ACE2, and antibodies, and C.L., and P.S. performed neutralization assays. P.S. isolated all Omicron variants. D.C., H.W., B.C., and N.T. provided materials. H.M.G. wrote mabscape and performed mapping and cluster analysis, including sequence and antigenic space analyses. A.J.M., D.S., T.G.R., A.A., S.B., S.A., S.A.J., P.K., E.B. S.J.D., A.J.P., T.L., and P.G. assisted with patient samples and vaccine trials. E.B., S.J.D., and P.K. conceived the study of vaccinated healthcare workers and oversaw the OPTIC Healthcare Worker study and sample collection/processing, G.R.S., and D.I.S. conceived the study and wrote the initial manuscript draft with other authors providing editorial comments. All authors read and approved the manuscript.

## Competing Financial Interests

G.R.S. sits on the GSK Vaccines Scientific Advisory Board and is a founder member of RQ Biotechnology. Oxford University holds intellectual property related to the Oxford-Astra Zeneca vaccine and SARS-CoV-2 mAb discovered in G.R.S’s laboratory. A.J.P. is Chair of UK Dept. Health and Social Care’s (DHSC) Joint Committee on Vaccination & Immunisation (JCVI) but does not participate in the JCVI COVID-19 committee, and is a member of the WHO’s SAGE. The views expressed in this article do not necessarily represent the views of DHSC, JCVI, or WHO. The University of Oxford has entered into a partnership with AstraZeneca on coronavirus vaccine development. T.L. is named as an inventor on a patent application covering this SARS-CoV-2 vaccine and was a consultant to Vaccitech for an unrelated project whilst the study was conducted. S.J.D. is a Scientific Advisor to the Scottish Parliament on COVID-19.

Figure S1 Neutralization curves for VH1-58 mAb. Pseudoviral neutralization curves for early pandemic mAb 253 (Dejnirattisai et al., 2021a) and Beta-47 (Liu et al., 2021b) against Victoria and the panel of Omicron lineage constructs.

Figure S2 Surface plasmon resonance (SPR) analysis of interaction between BA.2 or BA.4/5 RBD and selected mAbs. (A-F) Sensorgrams (Red: original binding curve; black: fitted curve) showing the interactions between BA.2 or BA.4/5 RBD and selected mAbs, with kinetics data shown. (G-K) Binding of BA.4/5 RBD is severely reduced compared to that of BA.2, so that the binding could not be accurately determined, as shown by a single-injection of 200 nM RBD over sample flow cells containing the mAb indicated.

## STAR Methods

### RESOURCE AVAILABILITY

#### Lead Contact

Resources, reagents and further information requirement should be forwarded to and will be responded by the Lead Contact, David I Stuart.

#### Materials Availability

Reagents generated in this study are available from the Lead Contact with a completed Materials Transfer Agreement.

### EXPERIMENTAL MODEL AND SUBJECT DETAILS

#### Bacterial Strains and Cell Culture

Vero (ATCC CCL-81) and VeroE6/TMPRSS2 cells were cultured at 37 °C in Dulbecco’s Modified Eagle medium (DMEM) high glucose (Sigma-Aldrich) supplemented with 10% fetal bovine serum (FBS), 2 mM GlutaMAX (Gibco, 35050061) and 100rnU/ml of penicillin– streptomycin. Human mAbs were expressed in HEK293T cells cultured in UltraDOMA PF Protein-free Medium (Cat# 12-727F, LONZA) at 37 °C with 5% CO_2_. HEK293T (ATCC CRL- 11268) cells were cultured in DMEM high glucose (Sigma-Aldrich) supplemented with 10% FBS, 1% 100X Mem Neaa (Gibco) and 1% 100X L-Glutamine (Gibco) at 37 °C with 5% CO_2_. To express RBD, RBD variants and ACE2, HEK293T cells were cultured in DMEM high glucose (Sigma) supplemented with 2% FBS, 1% 100X Mem Neaa and 1% 100X L-Glutamine at 37 °C for transfection. Omicron RBD and human mAbs were also expressed in HEK293T (ATCC CRL- 11268) cells cultured in FreeStyle 293 Expression Medium (ThermoFisher, 12338018) at 37 °C with 5% CO_2_. E.coli DH5α bacteria were used for transformation and large-scale preparation of plasmids. A single colony was picked and cultured in LB broth at 37 °C at 200 rpm in a shaker overnight.

#### Plasma from early pandemic and Alpha cases

Participants from the first wave of SARS-CoV2 in the U.K. and those sequence confirmed with B.1.1.7 lineage in December 2020 and February 2021 were recruited through three studies: Sepsis Immunomics [Oxford REC C, reference:19/SC/0296]), ISARIC/WHO Clinical Characterisation Protocol for Severe Emerging Infections [Oxford REC C, reference 13/SC/0149] and the Gastro-intestinal illness in Oxford: COVID sub study [Sheffield REC, reference: 16/YH/0247]. Diagnosis was confirmed through reporting of symptoms consistent with COVID-19 and a test positive for SARS-CoV-2 using reverse transcriptase polymerase chain reaction (RT-PCR) from an upper respiratory tract (nose/throat) swab tested in accredited laboratories. A blood sample was taken following consent at least 14 days after symptom onset. Clinical information including severity of disease (mild, severe or critical infection according to recommendations from the World Health Organisation) and times between symptom onset and sampling and age of participant was captured for all individuals at the time of sampling. Following heat inactivation of plasma/serum samples they were aliquoted so that no more than 3 freeze thaw cycles were performed for data generation.

#### Sera from Beta, Gamma and Delta and BA.1 infected cases

Beta and Delta samples from UK infected cases were collected under the “Innate and adaptive immunity against SARS-CoV-2 in healthcare worker family and household members” protocol affiliated to the Gastro-intestinal illness in Oxford: COVID sub study discussed above and approved by the University of Oxford Central University Research Ethics Committee. All individuals had sequence confirmed Beta/Delta infection or PCR-confirmed symptomatic disease occurring whilst in isolation and in direct contact with Beta/Delta sequence-confirmed cases. Additional Beta infected serum (sequence confirmed) was obtained from South Africa. At the time of swab collection patients signed an informed consent to consent for the collection of data and serial blood samples. The study was approved by the Human Research Ethics Committee of the University of the Witwatersrand (reference number 200313) and conducted in accordance with Good Clinical Practice guidelines. Gamma samples were provided by the International Reference Laboratory for Coronavirus at FIOCRUZ (WHO) as part of the national surveillance for coronavirus and had the approval of the FIOCRUZ ethical committee (CEP 4.128.241) to continuously receive and analyse samples of COVID-19 suspected cases for virological surveillance. Clinical samples were shared with Oxford University, UK under the MTA IOC FIOCRUZ 21-02.

#### Sera from BA.1 infected cases, study subjects

Following informed consent, individuals with omicron BA.1 were co-enrolled into the ISARIC/WHO Clinical Characterisation Protocol for Severe Emerging Infections [Oxford REC C, reference 13/SC/0149] and the “Innate and adaptive immunity against SARS-CoV-2 in healthcare worker family and household members” protocol affiliated to the Gastro- intestinal illness in Oxford: COVID sub study [Sheffield REC, reference: 16/YH/0247] further approved by the University of Oxford Central University Research Ethics Committee.

Diagnosis was confirmed through reporting of symptoms consistent with COVID-19 or a positive contact of a known Omicron case, and a test positive for SARS-CoV-2 using reverse transcriptase polymerase chain reaction (RT-PCR) from an upper respiratory tract (nose/throat) swab tested in accredited laboratories and lineage sequence confirmed through national reference laboratories. A blood sample was taken following consent at least 10 days after PCR test confirmation. Clinical information including severity of disease (mild, severe or critical infection according to recommendations from the World Health Organisation) and times between symptom onset and sampling and age of participant was captured for all individuals at the time of sampling.

#### Sera from Pfizer vaccinees

Pfizer vaccine serum was obtained from volunteers who had received three doses of the BNT162b2 vaccine. Vaccinees were Health Care Workers, based at Oxford University Hospitals NHS Foundation Trust, not known to have prior infection with SARS-CoV-2 and were enrolled in the OPTIC Study as part of the Oxford Translational Gastrointestinal Unit GI Biobank Study 16/YH/0247 [research ethics committee (REC) at Yorkshire & The Humber – Sheffield] which has been amended for this purpose on 8 June 2020. The study was conducted according to the principles of the Declaration of Helsinki (2008) and the International Conference on Harmonization (ICH) Good Clinical Practice (GCP) guidelines. Written informed consent was obtained for all participants enrolled in the study. Participants were sampled approximately 28 days (range 25-56) after receiving a third “booster dose of BNT162B2 vaccine. The mean age of vaccinees was 37 years (range 22-66), 21 male and 35 female.

#### AstraZeneca-Oxford vaccine study procedures and sample processing

Full details of the randomized controlled trial of ChAdOx1 nCoV-19 (AZD1222), were previously published (PMID: 33220855/PMID: 32702298). These studies were registered at ISRCTN (15281137 and 89951424) and ClinicalTrials.gov (NCT04324606 and NCT04400838). Written informed consent was obtained from all participants, and the trial is being done in accordance with the principles of the Declaration of Helsinki and Good Clinical Practice. The studies were sponsored by the University of Oxford (Oxford, UK) and approval obtained from a national ethics committee (South Central Berkshire Research Ethics Committee, reference 20/SC/0145 and 20/SC/0179) and a regulatory agency in the United Kingdom (the Medicines and Healthcare Products Regulatory Agency). An independent DSMB reviewed all interim safety reports. A copy of the protocols was included in previous publications (Folegatti et al., 2020). Data from vaccinated volunteers who received three vaccinations are included in this study. Blood samples were collected and serum separated approximately 28 days (range 26- 34 days) following the third dose.

### Method Details

#### Plasmid construction and pseudotyped lentiviral particles production

Pseudotyped lentivirus expressing SARS-CoV-2 S proteins from ancestral strain (Victoria, S247R), BA.1, BA.1.1, and BA.2 were constructed as described previously (Nie et al., 2020, Liu et al., 2021b, Nutalai et al., 2022), with some modifications. A similar strategy was applied for BA.3 and BA.4/5, briefly, BA.3 mutations were constructed using the combination fragments from BA.1 and BA.2. The resulting mutations are as follows, A67V, Δ69-70, T95I, G142D, Δ143-145, Δ211/L212I, G339D, S371F, S373P, S375F, D405N, K417N, N440K, G446S, S477N, T478K, E484A, Q493R, Q498R, N501Y, Y505H, D614G, H655Y, N679K, P681H, N764K, D796Y, Q954H, and N969K. Although BA.4/5 S protein shared some amino acid mutations with BA.2 (Nutalai et al., 2022), to generate BA.4/5 we added mutations Δ69-70, L452R, F486V, and R498Q. The resulting S gene-carrying pcDNA3.1 was used for generating pseudoviral particles together with the lentiviral packaging vector and transfer vector encoding luciferase reporter. Integrity of contructs was sequence confirmed.

#### Pseudoviral neutralization test

The details of the pseudoviral neutralization test are as described previously (Liu et al., 2021b) with some modifications. Briefly, the neutralizing activity of potent monoclonal antibodies generated from donors who had recovered from Omicron were assayed against Victoria, Omicron-BA.1, BA.1.1, BA.2, BA.3 and BA.4/5. Four-fold serial dilutions of each mAb were incubated with pseudoviral particles at 37°C, 5% CO2 for 1 hr. Stable HEK293T/17 cells expressing human ACE2 were then added to the mixture at 1.5 x 10^4^ cells/well. 48 hr post transduction, culture supernatants were removed and 50 µL of 1:2 Bright-GloTM Luciferase assay system (Promega, USA) in 1x PBS was added to each well. The reaction was incubated at room temperature for 5 mins and firefly luciferase activity was measured using CLARIOstar® (BMG Labtech, Ortenberg, Germany). The percentage neutralization was calculated relative to the control. Probit analysis was used to estimate the dilution that inhibited half maximum pseudotyped lentivirus infection (PVNT50).

To determine the neutralizing activity of convalescent plasma/serum samples or vaccine sera, 3-fold serial dilutions of samples were incubated with pseudoviral particles for 1 hr and the same strategy as mAb was applied.

#### Cloning of RBDs

To generate His-tagged constructs of BA.4/5 RBD, site-directed PCR mutagenesis was performed using the BA.2 RBD construct as the template (Nutalai et al., 2022), with the introduction of L452R, F486V and R493Q mutations. The gene fragment was amplified with pNeoRBD333Omi_F (5’- GGTTGCGTAGCTGAAACCGGTCATCACCATCACCATCACACCAATCTGTGCCCTTTCGAC-3’) and pNeoRBD333_R (5’-GTGATGGTGGTGCTTGGTACCTTATTACTTCTTGCCGCACACGGTAGC-3’), and cloned into the pNeo vector (Supasa et al., 2021). To generate the BA.4/5 RBD construct containing a BAP-His tag, the gene fragment was amplified with RBD333_F (5’- GCGTAGCTGAAACCGGCACCAATCTGTGCCCTTTCGAC-3’) and RBD333_BAP_R (5’-GTCATTCAGCAAGCTCTTCTTGCCGCACACGGTAGC-3’), and cloned into the pOPINTTGneo-BAP vector (Huo et al., 2020a). Cloning was performed using the ClonExpress II One Step Cloning Kit (Vazyme). The Constructs were verified by Sanger sequencing after plasmid isolation using QIAGEN Miniprep kit (QIAGEN).

#### Production of RBDs

Plasmids encoding RBDs were transfected into Expi293F™ Cells (ThermoFisher) by PEI, cultured in FreeStyle™ 293 Expression Medium (ThermoFisher) at 30 °C with 8% CO_2_ for 4 days. To express biotinylated RBDs, the RBD-BAP plasmid was co-transfected with pDisplay-BirA-ER (Addgene plasmid 20856; coding for an ER-localized biotin ligase), in the presence of 0.8 mM D-biotin (Sigma-Aldrich). The conditioned medium was diluted 1:2 into binding buffer (50 mM sodium phosphate, 500 mM sodium chloride, pH 8.0). RBDs were purified with a 5 mL HisTrap nickel column (GE Healthcare) through His-tag binding, followed by a Superdex 75 10/300 GL gel filtration column (GE Healthcare) in 10 mM HEPES and 150 mM sodium chloride.

#### Surface Plasmon Resonance

The surface plasmon resonance experiments were performed using a Biacore T200 (GE Healthcare). All assays were performed with running buffer of HBS-EP (Cytiva) at 25C°C.

To determine the binding kinetics between the RBDs and mAb Omi-32 / Omi-42, a Biotin CAPture Kit (Cytiva) was used. Biotinylated RBD was immobilised onto the sample flow cell of the sensor chip. The reference flow cell was left blank. The mAb Fab was injected over the two flow cells at a range of five concentrations prepared by serial two-fold dilutions, at a flow rate of 30CμlCmin using a single-cycle kinetics programme. Running buffer was also injected using the same programme for background subtraction. All data were fitted to a 1:1 binding model using Biacore T200 Evaluation Software 3.1.

To determine the binding kinetics between RBDs and ACE2 / other mAbs, a Protein A sensor chip (Cytiva) was used. ACE2-Fc or mAb in the IgG form was immobilised onto the sample flow cell of the sensor chip. The reference flow cell was left blank. RBD was injected over the two flow cells at a range of five concentrations prepared by serial two-fold dilutions, at a flow rate of 30CμlCmin using a single-cycle kinetics programme. Running buffer was also injected using the same programme for background subtraction. All data were fitted to a 1:1 binding model using Biacore T200 Evaluation Software 3.1.

To determine the binding affinity of BA.4/5 RBD and mAb Omi-12, a Protein A sensor chip (Cytiva) was used. The Ig Omi-12 was immobilised onto the sample flow cell of the sensor chip. The reference flow cell was left blank. RBD was injected over the two flow cells at a range of seven concentrations prepared by serial twofold dilutions, at a flow rate of 30CμlCmin . Running buffer was also injected using the same programme for background subtraction. All data were fitted to a 1:1 binding model using Prism9 (GraphPad).

To compare the binding profiles between BA.2 and BA.4/5 RBD for mAb Omi-06 / Omi-25 / Omi-26, a Protein A sensor chip (Cytiva) was used. mAb in the IgG form was immobilised onto the sample flow cell of the sensor chip to a similar level (∼350 RU). The reference flow cell was left blank. A single injection of RBD was performed over the two flow cells at 200 nM, at a flow rate of 30CμlCmin . Running buffer was also injected using the same programme for background subtraction. The sensorgrams were plotted using Prism9 (GraphPad).

To compare the binding profiles between BA.2 and BA.4/5 RBD for mAb Omi-02 / Omi-23 / Omi-31, a Biotin CAPture Kit (Cytiva) was used. Biotinylated BA.2 and BA.4/5 RBD was immobilised onto the sample flow cell of the sensor chip to a similar level (∼120 RU). The reference flow cell was left blank. A single injection of mAb Fab was performed over the two flow cells at 200 nM, at a flow rate of 30CμlCmin . Running buffer was also injected using the same programme for background subtraction. The sensorgrams were plotted using Prism9 (GraphPad).

#### IgG mAbs and Fabs production

AstraZeneca and Regeneron antibodies were provided by AstraZeneca, Vir, Lilly and Adagio antibodies were provided by Adagio. For the in-house antibodies, heavy and light chains of the indicated antibodies were transiently transfected into 293Y cells and antibody purified from supernatant on protein A as previously described (Nutalai et al., 2022). Fabs were digested from purified IgGs with papain using a Pierce Fab Preparation Kit (Thermo Fisher), following the manufacturer’s protocol.

### Antigenic mapping

Antigenic mapping of omicron was carried out through an extension of a previous algorithm (Liu et al., 2021a). In short, coronavirus variants were assigned three-dimensional coordinates whereby the distance between two points indicates the base drop in neutralization titre. Each serum was assigned a strength parameter which provided a scalar offset to the logarithm of the neutralization titre. These parameters were refined to match predicted neutralization titres to observed values by taking an average of superimposed positions from 30 separate runs. The three-dimensional positions of the variants of concern: Victoria, Alpha, Beta, Gamma, Delta and Omicron were plotted for display.

### Quantification and statistical analysis

Statistical analyses are reported in the results and figure legends. Neutralization was measured on pseudovirus. The percentage reduction was calculated and IC_50_ determined using the probit program from the SPSS package. The Wilcoxon matched-pairs signed rank test was used for the analysis and two-tailed P values were calculated on geometric mean values.

