## Supplementary Tables and Figure for "Further antibody escape by Omicron BA.4 and BA.5 from vaccine and BA.1 serum"

### Slide 1
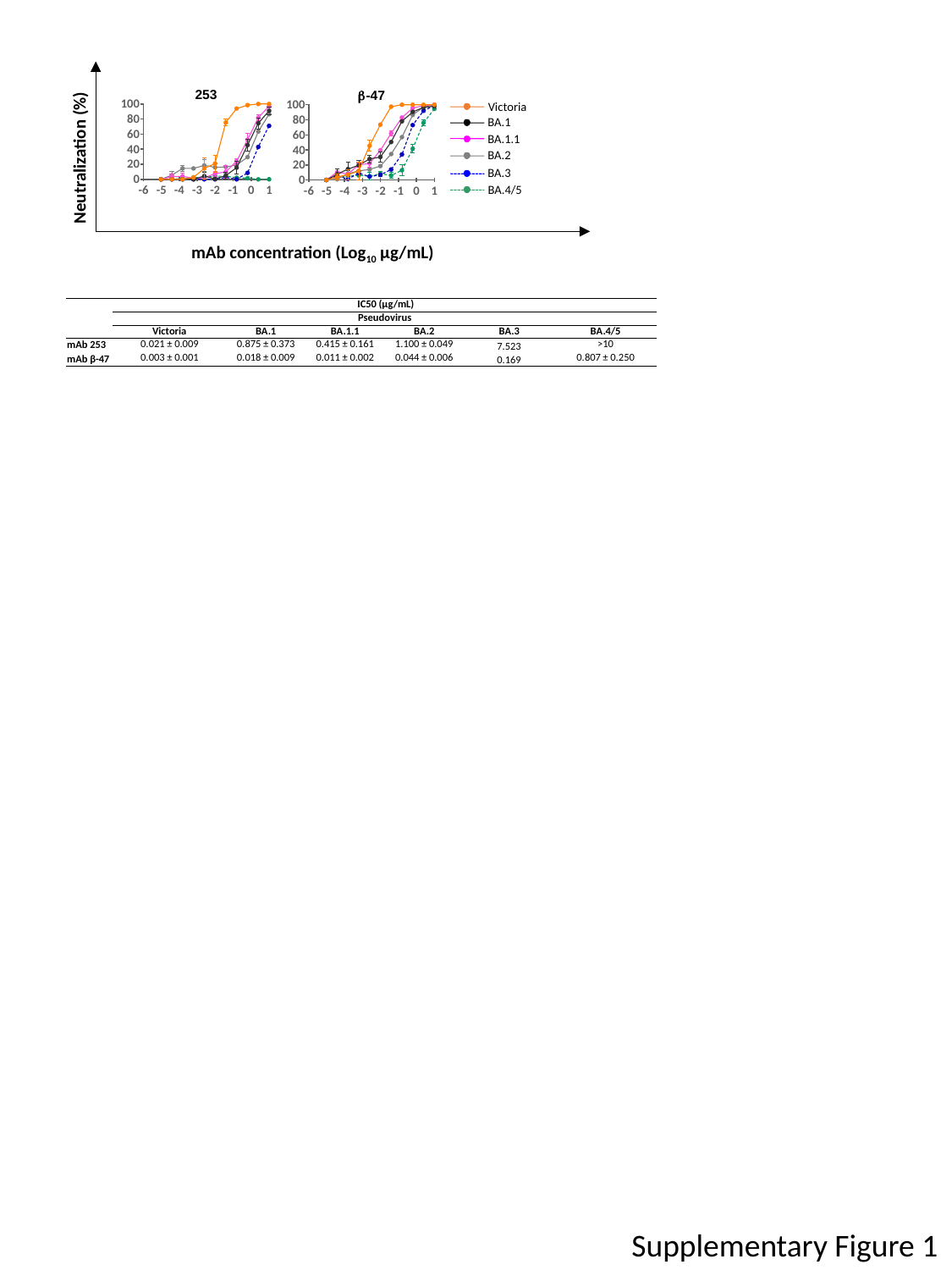

Victoria
BA.1
BA.1.1
BA.2
BA.3
BA.4/5
Neutralization (%)
mAb concentration (Log_10_ µg/mL)
| | IC50 (µg/mL) | | | | | |
| --- | --- | --- | --- | --- | --- | --- |
| | Pseudovirus | | | | | |
| | Victoria | BA.1 | BA.1.1 | BA.2 | BA.3 | BA.4/5 |
| mAb 253 | 0.021 ± 0.009 | 0.875 ± 0.373 | 0.415 ± 0.161 | 1.100 ± 0.049 | 7.523 | >10 |
| mAb β-47 | 0.003 ± 0.001 | 0.018 ± 0.009 | 0.011 ± 0.002 | 0.044 ± 0.006 | 0.169 | 0.807 ± 0.250 |
Supplementary Figure 1

### Slide 2
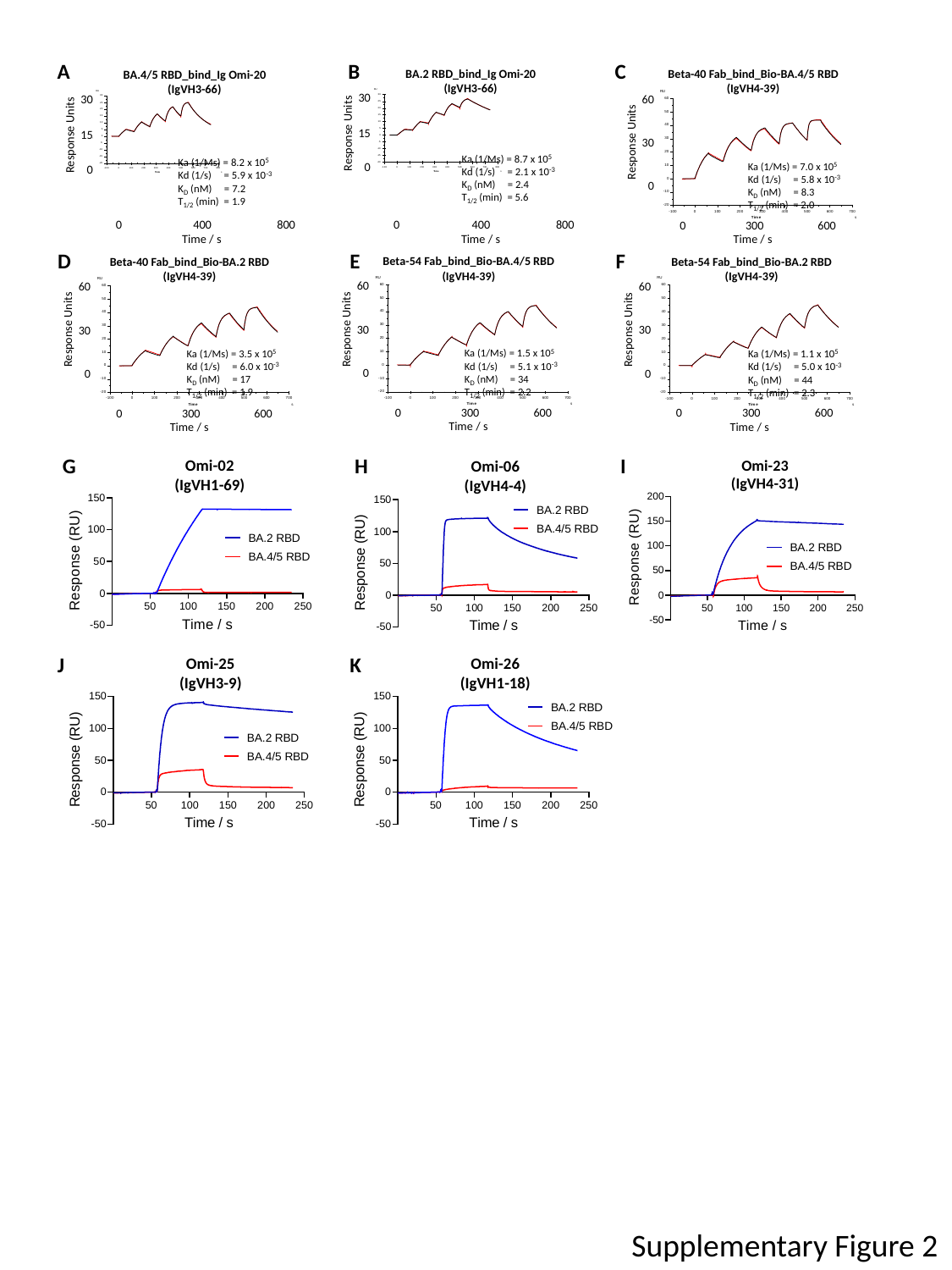

C
B
A
D
E
F
G
H
I
J
K
Supplementary Figure 2

### Slide 3
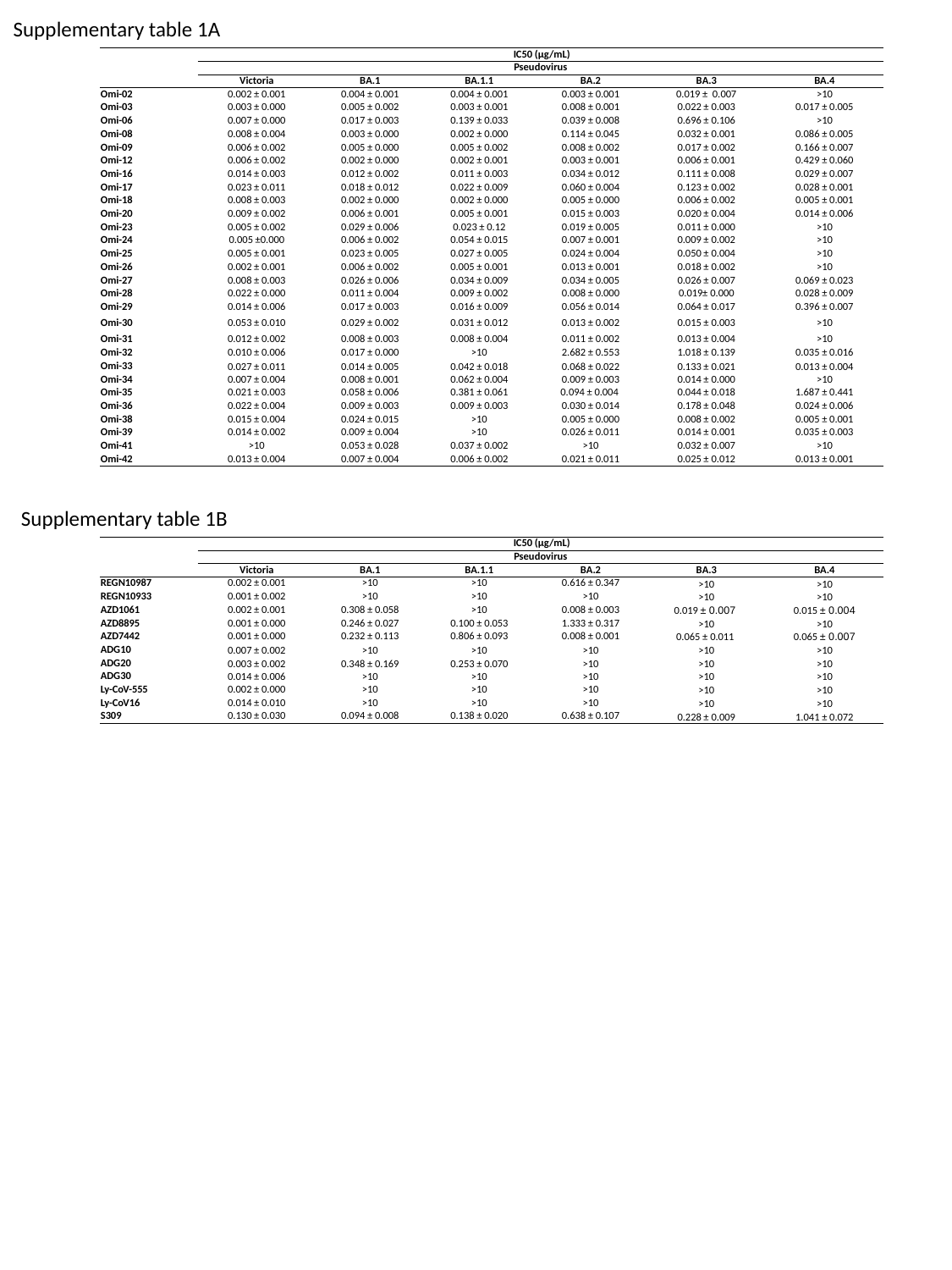

Supplementary table 1A
| | IC50 (µg/mL) | | | | | |
| --- | --- | --- | --- | --- | --- | --- |
| | Pseudovirus | | | | | |
| | Victoria | BA.1 | BA.1.1 | BA.2 | BA.3 | BA.4 |
| Omi-02 | 0.002 ± 0.001 | 0.004 ± 0.001 | 0.004 ± 0.001 | 0.003 ± 0.001 | 0.019 ± 0.007 | >10 |
| Οmi-03 | 0.003 ± 0.000 | 0.005 ± 0.002 | 0.003 ± 0.001 | 0.008 ± 0.001 | 0.022 ± 0.003 | 0.017 ± 0.005 |
| Οmi-06 | 0.007 ± 0.000 | 0.017 ± 0.003 | 0.139 ± 0.033 | 0.039 ± 0.008 | 0.696 ± 0.106 | >10 |
| Οmi-08 | 0.008 ± 0.004 | 0.003 ± 0.000 | 0.002 ± 0.000 | 0.114 ± 0.045 | 0.032 ± 0.001 | 0.086 ± 0.005 |
| Οmi-09 | 0.006 ± 0.002 | 0.005 ± 0.000 | 0.005 ± 0.002 | 0.008 ± 0.002 | 0.017 ± 0.002 | 0.166 ± 0.007 |
| Οmi-12 | 0.006 ± 0.002 | 0.002 ± 0.000 | 0.002 ± 0.001 | 0.003 ± 0.001 | 0.006 ± 0.001 | 0.429 ± 0.060 |
| Οmi-16 | 0.014 ± 0.003 | 0.012 ± 0.002 | 0.011 ± 0.003 | 0.034 ± 0.012 | 0.111 ± 0.008 | 0.029 ± 0.007 |
| Οmi-17 | 0.023 ± 0.011 | 0.018 ± 0.012 | 0.022 ± 0.009 | 0.060 ± 0.004 | 0.123 ± 0.002 | 0.028 ± 0.001 |
| Οmi-18 | 0.008 ± 0.003 | 0.002 ± 0.000 | 0.002 ± 0.000 | 0.005 ± 0.000 | 0.006 ± 0.002 | 0.005 ± 0.001 |
| Οmi-20 | 0.009 ± 0.002 | 0.006 ± 0.001 | 0.005 ± 0.001 | 0.015 ± 0.003 | 0.020 ± 0.004 | 0.014 ± 0.006 |
| Οmi-23 | 0.005 ± 0.002 | 0.029 ± 0.006 | 0.023 ± 0.12 | 0.019 ± 0.005 | 0.011 ± 0.000 | >10 |
| Οmi-24 | 0.005 ±0.000 | 0.006 ± 0.002 | 0.054 ± 0.015 | 0.007 ± 0.001 | 0.009 ± 0.002 | >10 |
| Οmi-25 | 0.005 ± 0.001 | 0.023 ± 0.005 | 0.027 ± 0.005 | 0.024 ± 0.004 | 0.050 ± 0.004 | >10 |
| Οmi-26 | 0.002 ± 0.001 | 0.006 ± 0.002 | 0.005 ± 0.001 | 0.013 ± 0.001 | 0.018 ± 0.002 | >10 |
| Οmi-27 | 0.008 ± 0.003 | 0.026 ± 0.006 | 0.034 ± 0.009 | 0.034 ± 0.005 | 0.026 ± 0.007 | 0.069 ± 0.023 |
| Οmi-28 | 0.022 ± 0.000 | 0.011 ± 0.004 | 0.009 ± 0.002 | 0.008 ± 0.000 | 0.019± 0.000 | 0.028 ± 0.009 |
| Οmi-29 | 0.014 ± 0.006 | 0.017 ± 0.003 | 0.016 ± 0.009 | 0.056 ± 0.014 | 0.064 ± 0.017 | 0.396 ± 0.007 |
| Οmi-30 | 0.053 ± 0.010 | 0.029 ± 0.002 | 0.031 ± 0.012 | 0.013 ± 0.002 | 0.015 ± 0.003 | >10 |
| Οmi-31 | 0.012 ± 0.002 | 0.008 ± 0.003 | 0.008 ± 0.004 | 0.011 ± 0.002 | 0.013 ± 0.004 | >10 |
| Οmi-32 | 0.010 ± 0.006 | 0.017 ± 0.000 | >10 | 2.682 ± 0.553 | 1.018 ± 0.139 | 0.035 ± 0.016 |
| Οmi-33 | 0.027 ± 0.011 | 0.014 ± 0.005 | 0.042 ± 0.018 | 0.068 ± 0.022 | 0.133 ± 0.021 | 0.013 ± 0.004 |
| Οmi-34 | 0.007 ± 0.004 | 0.008 ± 0.001 | 0.062 ± 0.004 | 0.009 ± 0.003 | 0.014 ± 0.000 | >10 |
| Οmi-35 | 0.021 ± 0.003 | 0.058 ± 0.006 | 0.381 ± 0.061 | 0.094 ± 0.004 | 0.044 ± 0.018 | 1.687 ± 0.441 |
| Οmi-36 | 0.022 ± 0.004 | 0.009 ± 0.003 | 0.009 ± 0.003 | 0.030 ± 0.014 | 0.178 ± 0.048 | 0.024 ± 0.006 |
| Οmi-38 | 0.015 ± 0.004 | 0.024 ± 0.015 | >10 | 0.005 ± 0.000 | 0.008 ± 0.002 | 0.005 ± 0.001 |
| Οmi-39 | 0.014 ± 0.002 | 0.009 ± 0.004 | >10 | 0.026 ± 0.011 | 0.014 ± 0.001 | 0.035 ± 0.003 |
| Οmi-41 | >10 | 0.053 ± 0.028 | 0.037 ± 0.002 | >10 | 0.032 ± 0.007 | >10 |
| Οmi-42 | 0.013 ± 0.004 | 0.007 ± 0.004 | 0.006 ± 0.002 | 0.021 ± 0.011 | 0.025 ± 0.012 | 0.013 ± 0.001 |
Supplementary table 1B
| | IC50 (µg/mL) | | | | | |
| --- | --- | --- | --- | --- | --- | --- |
| | Pseudovirus | | | | | |
| | Victoria | BA.1 | BA.1.1 | BA.2 | BA.3 | BA.4 |
| REGN10987 | 0.002 ± 0.001 | >10 | >10 | 0.616 ± 0.347 | >10 | >10 |
| REGN10933 | 0.001 ± 0.002 | >10 | >10 | >10 | >10 | >10 |
| AZD1061 | 0.002 ± 0.001 | 0.308 ± 0.058 | >10 | 0.008 ± 0.003 | 0.019 ± 0.007 | 0.015 ± 0.004 |
| AZD8895 | 0.001 ± 0.000 | 0.246 ± 0.027 | 0.100 ± 0.053 | 1.333 ± 0.317 | >10 | >10 |
| AZD7442 | 0.001 ± 0.000 | 0.232 ± 0.113 | 0.806 ± 0.093 | 0.008 ± 0.001 | 0.065 ± 0.011 | 0.065 ± 0.007 |
| ADG10 | 0.007 ± 0.002 | >10 | >10 | >10 | >10 | >10 |
| ADG20 | 0.003 ± 0.002 | 0.348 ± 0.169 | 0.253 ± 0.070 | >10 | >10 | >10 |
| ADG30 | 0.014 ± 0.006 | >10 | >10 | >10 | >10 | >10 |
| Ly-CoV-555 | 0.002 ± 0.000 | >10 | >10 | >10 | >10 | >10 |
| Ly-CoV16 | 0.014 ± 0.010 | >10 | >10 | >10 | >10 | >10 |
| S309 | 0.130 ± 0.030 | 0.094 ± 0.008 | 0.138 ± 0.020 | 0.638 ± 0.107 | 0.228 ± 0.009 | 1.041 ± 0.072 |
